# Novel Biosensor Identifies Ruxolitinib as a Potent and Cardioprotective CaMKII Inhibitor

**DOI:** 10.1101/2022.09.24.509320

**Authors:** Oscar E. Reyes Gaido, Jonathan M. Granger, Lubika J. Nkashama, Brian L. Lin, Alan Long, Olurotimi O. Mesubi, Kate L. Schole, Chantelle E. Terrilion, Jun O. Liu, Elizabeth D. Luczak, Mark E. Anderson

## Abstract

Ca^2+^/Calmodulin-dependent protein kinase II (CaMKII) hyperactivity causes heart injury and arrhythmias—two major sources of mortality worldwide. Despite proven benefits of CaMKII inhibition in numerous preclinical models of heart disease, translation of CaMKII antagonists into humans has been stymied by low potency, toxicity, and an enduring concern for adverse effects on cognition due to an established role of CaMKII in learning and memory. To address these challenges, we asked if any clinically approved drugs, developed for other purposes, were potent CaMKII inhibitors. For this, we engineered a novel fluorescent biosensor, CaMKAR (CaMKII Activity Reporter), which features superior sensitivity, kinetics, and tractability for high throughput screening. Using this tool, we carried a drug repurposing screen (4,475 compounds in clinical use) in human cells expressing autonomously active CaMKII. This yielded five previously unrecognized CaMKII inhibitors with clinically relevant potency: ruxolitinib, baricitinib, silmitasertib, crenolanib, and abemaciclib. Standout among these, ruxolitinib, an orally bioavailable and U.S Food and Drug Administration (FDA)-approved medication, inhibited CaMKII in cultured cardiomyocytes and in mice at concentrations equivalent to human doses. 10-minute treatment in mice was sufficient to prevent atrial fibrillation— the most common clinical arrhythmia. At cardioprotective doses, ruxolitinib-treated mice behaved normally in established cognitive assays. Our results suggest that human CaMKII inhibition is feasible and safe, and support prompt clinical investigation of ruxolitinib for cardiac indications.

**One Sentence Summary:** We developed a CaMKII biosensor suitable for high throughput screening and identified ruxolitinib as a CaMKII inhibitor capable of rescuing cardiac arrhythmia.

## INTRODUCTION

Cardiovascular diseases are leading causes of morbidity and premature death, with nearly half a billion patients affected worldwide *(1, 2)*. The multifunctional Ca^2+^/calmodulin-dependent protein kinase II (CaMKII) contributes to physiological regulation of intracellular Ca^2+^ cycling in cardiomyocytes, but, surprisingly, is dispensable for normal cardiac function *(3–5)*. In contrast, excessive CaMKII activity is cardiotoxic and CaMKII inhibition is cardioprotective in multiple preclinical models of genetic and acquired heart disease *(6–15)*. Excessive myocardial CaMKII activity is, in part, a consequence of catecholamine stimulation but clinically approved drugs that inhibit β-adrenergic receptors (‘β blockers’) are ineffective at preventing increased CaMKII activity in myocardium obtained from heart failure patients *(16)*. Thus, developing safe and effective direct CaMKII inhibitor drugs is a major unmet need and translational goal.

Although several inhibitory modalities have been described, none have reached the clinic due to various limitations. KN-62 and KN-93 were the first CaMKII inhibitors *(17, 18)*. These allosteric inhibitors alleviate CaMKII-toxicity in a numerous animal models *(6, 13, 19, 20)*. While remaining popular as tool compounds, these were clinically hampered by low potency (IC_50_ >2 µM), prohibitive off-target toxicity to cardiac ion channels, and most importantly, they fail to inhibit autonomously hyperactive CaMKII *(21–23)*. In fact, KN-93 has been recently shown to be a calmodulin inhibitor, which explains its broad binding profile and inability to inhibit active CaMKII *(24, 25)*. CaMKII inhibitory peptides (CaMKII-IN, AIP, CN19o) have served as powerful genetic research tools with impressive affinity and specificity *(26–28)*; yet these have failed to translate due to the challenges of peptide delivery: short plasma half-life, cell impermeability, and suboptimal viral delivery to human myocardium *(29)*. Aiming to address these limitations, several ATP-competitor small molecules have been identified or developed *(30–34)*. From these, only the natural product derivative DiOHF has made it to a human trial *(35)*; this study intended to inhibit CaMKII after ischemia/reperfusion injury but yielded negative results—thought to be due to low potency against CaMKII *(36)*. A major potential obstacle to developing CaMKII inhibitor therapies has been concern for toxicity related to learning and memory *(29)*. CaMKII is important for long-term potentiation and comprises a high fraction of the total protein in excitatory synapses *(37)*. We reasoned that discovery of an approved medication, preferably in broad use, with potent CaMKII inhibitory properties would provide critical insights to inform translational researchers, patients, and industry about the viability of CaMKII inhibitors for human use.

Development of small molecule inhibitors could be facilitated by improved biosensors with high throughput capability. A seminal biosensor, Camui, is composed of CaMKII fused to Förster resonance energy transfer (FRET) probes *(38)*. Camui reports CaMKII conformational changes as a proxy for activity. Our group previously developed CaMKII-KTR (Kinase Translocation Reporter) *(39)*. This reporter does detect CaMKII dependent target phosphorylation, but requires nucleocytoplasmic transport for its function, making it inadequate for high throughput applications. Based on these limitations, we sought to develop a CaMKII activity reporter with suitable properties for in cellulo screening.

In the present study, we address two major gaps outlined above. First, we engineered and validated a new genetically-encoded CaMKII activity biosensor for live cell and in vitro screening with high sensitivity. Next, we screened the safe-for-human pharmacopeia (4,475 compounds, see Methods) and identified several compounds capable of inhibiting hyperactive, autonomous CaMKII. From these, ruxolitinib displayed outstanding repurposing characteristics, including high potency, low cardiomyocyte toxicity, and a low brain:plasma concentration ratio. Ruxolitinib rescued atrial fibrillation in validated mouse model without impairing cognitive function.

## RESULTS

### Development of a CaMKII biosensor suitable for high throughput screening

Unimolecular fluorescent sensors have improved our ability to observe biological processes with spatiotemporal precision. The discovery of circularly permuted green fluorescent protein (cpGFP) has led to multiple sensors that detect ions, metabolites, and enzymes by coupling reconstitution of GFP fluorescence with the activity or presence of the molecule of interest *(40–42)*. Using this principle, we engineered kinase sensing cpGFP to detect CaMKII. We screened for peptides that rendered cpGFP sensitive to active CaMKII (Fig. S1A). Among sequences known to be CaMKII substrates, the CaMKII autophosphorylation peptide MHRQETVDCLK (amino acids 281-291 from human CAMKIIδ) led to the highest dynamic range (Fig. S1A). We refer to this sensor as CaMKAR (<u>CaMK</u>II <u>A</u>ctivity <u>R</u>eporter), composed of the CaMKII autophosphorylation peptide, cpGFP, and a phosphorylated amino acid binding domain from bacterial FHA1 (Fig. 1A). When CaMKII phosphorylates the linked substrate, the FHA1 domain binds to the phosphorylated peptide, causing a conformational change that allows reconstitution of the GFP chromophore. Akin to previous sensors, this reconstitution is excitation ratiometric: GFP emission (~520 nm light) excited by ~488 nm light increases whereas GFP emission excited by ~405 nm is unchanged or decreases. we therefore express CaMKAR signal as the ratio (R) of these two channels (Fig. 1A). To test CaMKAR dynamics, we treated CaMKAR-expressing HEK293T cells with ionomycin, causing intracellular Ca^2+^ influx and CaMKII activation (Fig. 1B and C). This resulted in a rapid increase in R relative to baseline (R/R_0_). Maximal stimulation of sensor with both ionomycin and constitutively active CaMKII overexpression revealed an in cellulo maximal dynamic range of 3.27 fold, or 227 ± 11.1% (Fig. S1B). Since the 400 nm channel is largely unchanged upon stimulation, CaMKAR can be used intensiometrically by quantifying the 488/520 nm channel intensity (Fig. S1C). This modality retains 96.4 ± 1.6% of the dynamic range compared to ratiometric mode.

**Figure 1.**
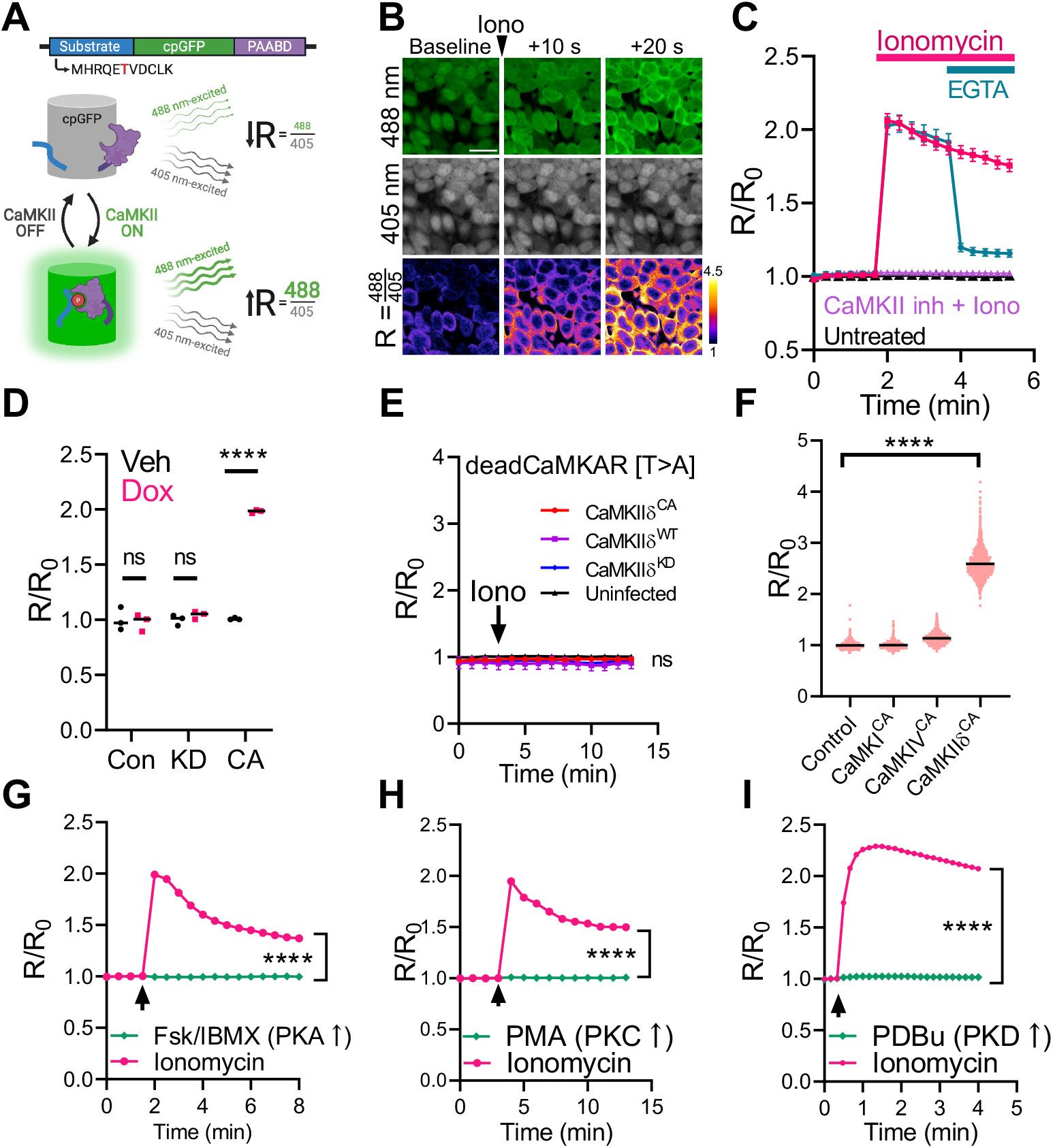
CaMKAR is a sensitive and specific CaMKII activity biosensor. **(A)** A schematic depiction of <u>CaMK</u>II <u>A</u>ctivity <u>R</u>eporter (CaMKAR) – (cpGFP: circularly permutated green fluorescent protein; PAABD: phosphorylated amino acid binding domain); see text for details. The phosphosubstrate threonine is shown in red. **(B)** Fluorescence confocal microscopy of CaMKAR-expressing 293T cells treated with ionomycin (5 μM). R= ex488/ex405. Scale bar = 20 μm. Third row is pseudo-colored to display fold-change over baseline. **(C)** Summary data for CaMKAR signal in HEK293T cells treated with vehicle (n=2,488-2,615 cells), ionomycin (5 μM; n=1,733-2,220 cells); pre-treatment with the CaMKII inhibitor (Inh) AS100397 (10 μM; n=1,305-1,348 cells) or post-treatment with EGTA (5 mM; n=1,474-1,746). R/R_0_ = R normalized to mean R prior to stimulation. (**D)** CaMKAR signal in HEK293T cells expressing doxycycline-induced wild type (n=3 wells) kinase-dead (KD; n=3 wells) or constitutively active (CA; n=3 wells) CaMKII. **(E)** DeadCaMKAR (CaMKAR_T6A_)-expressing HEK293T cells infected with wild-type (WT; n=mean of 3 wells), constitutively active (CA, n=mean of 3 wells), kinase dead (KD; n=mean of 3 wells) CaMKIIδ_C_ or uninfected (n=mean of 3 wells) and stimulated with ionomycin. **(F)** CaMKAR-expressing HEK293T cells co-transfected with constitutively active (CA) CaMKI (n=1,471 cells), CaMKIV (n=1,732 cells), or CaMKIIδ (n=1,816 cells). **(G)** CaMKAR-expressing 293T cells treated with Fsk (50 μM)/IBMX (100 μM; n=1,834-1891 cells) or ionomycin (n=1,997-2,293 cells). **(H)** CaMKAR-expressing 293T cells treated with PMA (100 ng/mL; n=696-737 cells), ionomycin (5 μM; n=767-916), or vehicle (see Methods). **(I)** CaMKAR-expressing 293T cells treated with PDBu (200 nM; n=1706-1736) or ionomycin (5 μM; n=1206-1228). Data shown as mean ± SEM (unless too small to display). Arrows denote treatment start. All observations done in biological triplicate. ns=p>0.05, ****p<0.0001; significance was determined via two-way ANOVA and Sidak’s multiple comparisons test (*D, G-I*); linear regression (*E*); one-way ANOVA with Dunnett’s multiple comparisons test (*F*).

Pretreatment with the tool CaMKII-inhibitor AS100397 *(43, 44)* eliminates and post-treatment with the Ca^2+^-chelator EGTA rapidly reduces CaMKAR signal (Fig. 1C). We then showed that CaMKAR sensing occurs via *bona fide* CaMKII-catalyzed phosphorylation by 3 orthogonal approaches: *1)* expression of constitutively active CaMKII^T287D^, but not kinase-defective CaMKII^K43M^, increases CaMKAR signal (Fig. 1D); *2)* CaMKAR sensing to both genetic and pharmacological activation of CaMKII is completely abolished when the substrate threonine residue is mutated to alanine (Fig. 1E); and *3)* CaMKAR signal is enhanced and fails to resolve when co-incubated with the phosphatase inhibitor Calyculin A (Fig. S1D). CaMKAR is sensitive to all 4 human CaMKII isoforms (α, β, γ, δ) and all 3 prevalent splice variants of CaMKIIδ (δC, δB, δ9) (Fig. S1E and F).

Next, we sought to benchmark CaMKAR’s performance and specificity. Camui was the first CaMKII biosensor developed and is still widely used *(38, 45)*. However, it has several limitations. Camui inherently contains and overexpresses CaMKIIα, a major brain isoform, has relatively low dynamic range (~5-70% vs CaMKAR’s 227 ± 11.1%), and reports conformational change rather than actual activity. In HEK293T cells, CaMKAR has nearly 10-fold greater signal-to-noise (Fig. S1G) and is ~3-fold faster (τ_CaMKAR_ =7.4 vs τ_Camui_=21.6) (Fig. S1H). Thus, CaMKAR enjoys improved sensitivity and kinetics.

We then asked if CaMKAR was exclusively reporting on CaMKII activity. Ca^2+^ mobilizing agents such as ionomycin can potentially activate many kinases. We set to determine whether CaMKAR is sensitive to other closely related and Ca^2+^-responsive kinases. CaMKAR is insensitive to constitutively active CaMKI and CaMKIV (Fig. 1F). CaMKAR also failed to sense Forskolin/IBMX-mediated PKA activation, PMA-mediated PKC activation, and PDBu-mediated PKD activation (Fig. 1G-I). We verified that the concentrations of Forskolin/IBMX and PMA used in these studies were sufficient to elicit PKA (Fig. S1I) and PKC activity (Fig. S1J). Lastly, CaMKAR’s response to ionomycin was unhindered by Gö6976, a dual PKC/PKD inhibitor (Fig. S1K).

Given its unimolecular architecture, we sought to leverage CaMKAR in an in vitro assay. We found that recombinant CaMKAR is fully functional and exhibits appropriate spectral changes upon incubation with CaMKII (Fig. S2A). These changes are ATP-dependent, further confirming their dependence on phosphorylation (Fig. S2A-D). CaMKAR displays a somewhat larger dynamic range in vitro (264 ± 2.1%) compared to cell-based assays (227 ± 11.1%), mostly due to a decrease in the 405 nm-excited emission when it is phosphorylated. Notably, the rate of the reaction is determined by the amount of CaMKII (Fig. S2E). Altogether, we have demonstrated that CaMKAR is a *bona fide* CaMKII activity biosensor with unprecedented sensitivity, fast kinetics, specificity, and suitability for live cell and in vitro drug screening.

### CaMKAR-based screen of drugs in clinical use

We hypothesized that among the clinically-approved pharmacopeia, we could find drugs that are both safe for human consumption and potent CaMKII inhibitors. Due to its high throughput tractability, CaMKAR is uniquely configured to answer this question. For screening, we first created a stable line of human K562 cells that co-express CaMKAR and doxycycline-inducible CaMKIIδ^T287D^: K562^CaMKII-CaMKAR^ (Fig. 2A). We chose K562 cells because they grow at high density and remain viable after expression of constitutively active CaMKII (Fig. S3A). Importantly, the use of constitutively active CaMKII instead of Ca^2+^-activated CaMKII increases the likelihood of identifying catalytic inhibitors rather than calmodulin inhibitors (such as KN-93). The sensitivity of CaMKAR enables screening to occur in small culture volumes. We found that CaMKAR signal inhibition by AS100397 is detectable in 30 μL of cell culture (Fig S3B). In our primary screen, we tested this cell line against the Johns Hopkins Drug Library v3.0— constructed by pooling 4,475 compounds approved for human use by regulatory agencies from the U.S., Europe, Japan, and China. This library targets over 70 different pathways and contains >130 known kinase inhibitors (data file S1). After 12 hours of treatment, cells were assayed for CaMKAR signal by high content imaging. Among drugs with an inhibitory signal, we found 118 compounds that reduced CaMKII activity by 60% or more, representing a false discovery-adjusted p value cutoff of < 3 × 10^−5^ (Fig. 2B; data file S1). We reasoned that these hits likely contained a mixture of genuine inhibitors, auto-fluorescent compounds, and phosphatase activators. To discern *bona fide* inhibitors from false positives, we performed a secondary screen using our CaMKAR in vitro assay. In this assay baseline fluorescence was measured in a mixture of CaMKII, Ca^2+^-bound calmodulin (Ca^2+^/CaM), and CaMKAR. Fluorescence was then measured after addition of drug, and then CaMKII catalysis was initiated by ATP. Thus, this assay simultaneously determines if a drug is auto-fluorescent and/or a direct inhibitor. This revealed 13 compounds with statistically significant inhibition (Fig. 2C, data file S2). Importantly, both screening steps returned the positive control, pan-kinase inhibitor staurosporine. To filter out weak inhibitors and further validate our results, these compounds were re-tested in HEK293T cells by us (Fig. S4) and in vitro by an independent commercial laboratory (data file S2). These steps confirmed 5 potent CaMKII inhibitors: ruxolitinib, crenolanib, baricitinib, abemaciclib, and silmitasertib (Figs. 2D, S5), all known ATP-competitors. Surprisingly, none of the compounds were designed against kinases in the CAMK superfamily, and 3 out of the 5 compounds were originally intended against tyrosine kinases—one of the most dissimilar kinase families relative to CaMK, as represented by kinase domain homology (Fig 2D). Unlike KN-93 *(27)*, all 5 compounds inhibited Ca^2+^-independent, autonomously active CaMKII in HEK293T cells (Fig. 2E, left panel). In this assay, crenolanib, ruxolitinib, and baricitinib were more potent than AS100397, and all drugs except crenolanib were less toxic after a 12 hours exposure (Fig. 2E, right panel). Thus, ruxolitinib and baricitinib stand out as being more potent and less toxic than tool inhibitors in cultured cells.

**Figure 2.**
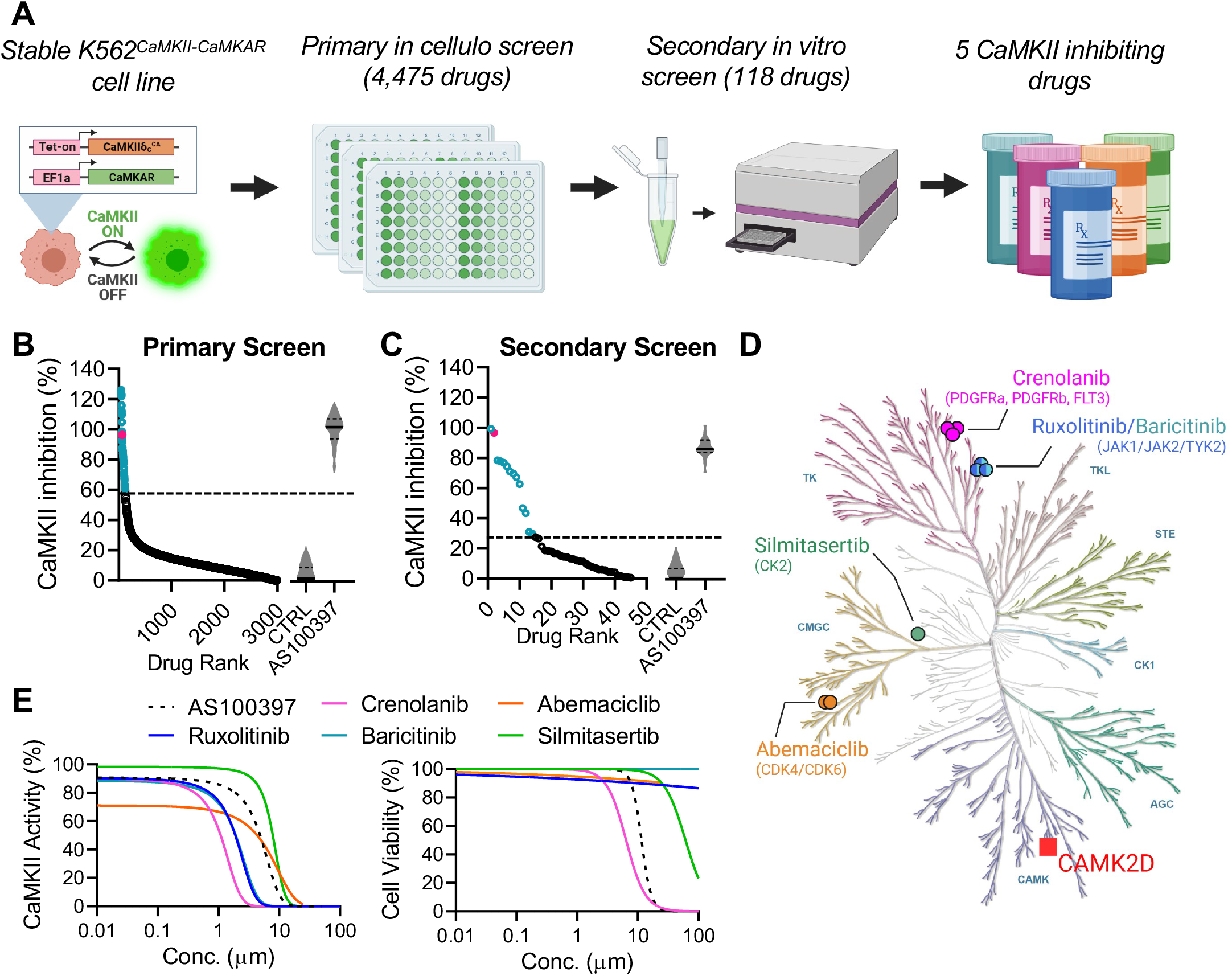
CaMKAR-based screen identifies CaMKII inhibitors amongst drugs in clinical use. **(A)** A schematic depiction of CaMKAR-based high throughput screen pipeline. K562 cells were co-infected with CaMKAR- and Doxycycline-inducible CaMKII_T287D_-encoding lentiviruses. K562_CaMKII-CaMKAR_ cells were screened against the Johns Hopkins Drug Library v3.0 in cellulo and subsequently in vitro (see Methods). **(B)** Drugs ranked according to in cellulo CaMKII inhibition in primary screen. CaMKII inhibition % defined by min-max normalization using means of control groups (*right; same data as fig. S3B*). 118 selected hits shown in blue. Positive control staurosporine shown in violet. Dashed line represents hit selection threshold (see Methods). Data shown are subset from complete data set (see Supplemental Materials). **(C)** Hits from *B* ranked according to in vitro CaMKII inhibition as detected by CaMKAR secondary screen. CaMKII inhibition % defined by max-normalization against untreated control (see methods). 13 identified hits in blue. Data shown are subset from complete data set (see Supplemental Materials). **(D)** Five CaMKII inhibitory drugs and their intended targets in the human kinase homology dendrogram. Kinase groups: AGC=Containing PKA, PKG, PKC families, CAMK=Calcium/calmodulin-dependent protein kinase, CK1=Casein Kinase 1, CMGC=Containing CDK, MAPK, GSK3, CLK, STE=Homologs of yeast Sterile 7, Sterile 11, Sterile 20, TK=Tyrosine kinase, TKL=Tyrosine kinase-like. **(E)** IC_50_ (*left*) and cell viability (*right*) curves from 293T cells expressing CaMKAR and CaMKII_T287D_ and exposed to drugs in *E* and tool inhibitor (AS100397, 10 mM). Measurements in *E* done in biological triplicate, complete data set shown in fig. S4.

### Clinically approved drugs inhibit CaMKII in cardiac cells

Because of the established benefits of CaMKII inhibition in cardiovascular disease models, we asked if these 5 compounds could inhibit CaMKII in cardiomyocytes. In neonatal rat ventricular myocytes (NRVMs), we are able to stimulate CaMKII activity by rapid field pacing (Fig. 3A and B). All five compounds inhibited this response at high doses (Fig. 3C). However, to determine which compounds are likely to have an effect at clinically achievable doses, we adjusted their concentration to their respective maximal human plasma concentration *(46–50)*. This revealed that ruxolitinib, crenolanib, abemaciclib, and silmitasertib, but not baricitinib, retained inhibition (Fig. 3D and E). Due to its high potency, low toxicity, and best-known safety profile among our hits, we henceforth decided to focus on ruxolitinib. Using this cardiomyocyte pacing assay, we found that ruxolitinib is over 10-fold more potent than 3’,4’-dihydroxyflavonol (DiOHF), a low-potency CaMKII inhibitor recently trialed in humans (Fig. 3F) *(34, 35)*. This inhibitory effect appears to be independent of ruxolitinib’s JAK1/2 inhibition. For one, we found multiple JAK1/2 inhibitors in our screen that failed to reduce CaMKAR signal (Fig. S6A). Among these compounds, their known IC_50_ values against JAK1/2 were not significantly correlated with CaMKAR inhibition in our screen (Fig. S6B). Furthermore, filgotinib, another FDA-approved compound with similar potency against JAK1/2 failed to inhibit pacing-induced CaMKII activity (Fig. S6C). 48-hour exposure in NRVMs revealed that ruxolitinib is well tolerated up to 100 μM. This was similar to DiOHF and vastly superior to other CaMKII inhibitors KN-93, AS100397, and Hesperadin (Fig. 3G). We conclude that the drugs identified in our screen shared the ability to inhibit CaMKII in cardiac cells. But among these, ruxolitinib appears to be best suited for repurposing due to its potency relative to its human dose and low toxicity.

**Figure 3.**
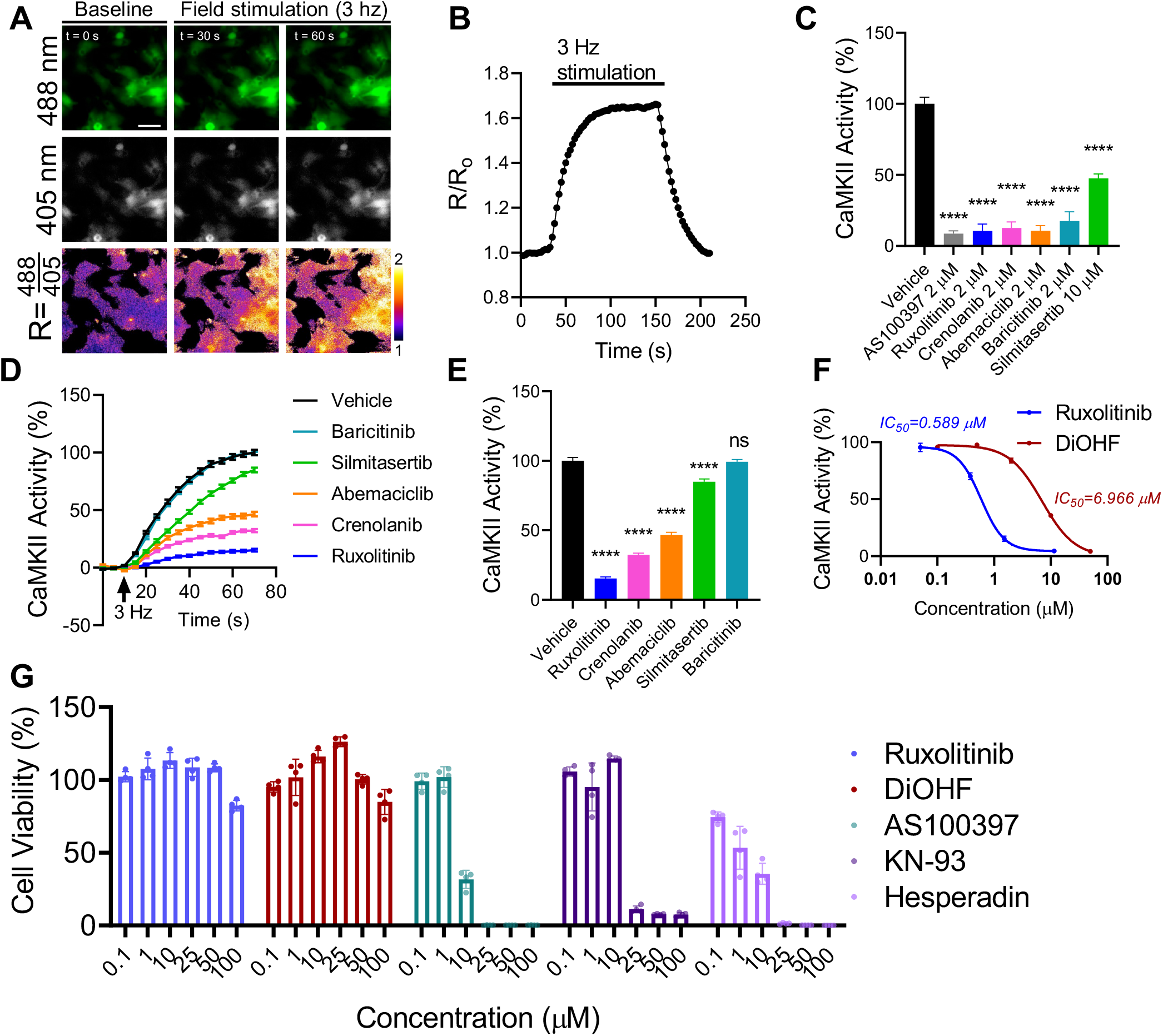
Drugs in clinical use inhibit CaMKII in cardiac cells. **(A)** Fluorescence imaging timelapse of CaMKAR-expressing neonatal rat ventricular cardiomyocytes (NRVMs) and **(B)** summary data during 3 Hz field pacing (n=1,065-1,160). R/R_0_=R normalized to mean R prior to stimulation. Scale bar = 50 μm. **(C)** Effect of treatment with vehicle (n=317), AS100397 (n=151) and FDA-approved drugs ruxolitinib (n=148), crenolanib (n=156), abemaciclib (n=138), baricitinib (n=136), and silmitasertib (n=149) on pacing-induced CaMKII activity in NRVMs. CaMKII inhibition % defined by min-max normalization between untreated R and maximally observed R. **(D)** CaMKII inhibition in NRVMs treated with drugs adjusted to their maximum human plasma concentration during 3 Hz pacing: ruxolitinib (1.51 μM, n=384), crenolanib (478 nM, n=302), abemaciclib (243 nM, n=302), baricitinib (58 nM n=334), and silmitasertib (3.42 μM, n=285). **(E)** Summary data from *D* at 60 seconds post stimulation. **(F)** IC_50_ curves for ruxolitinib (n=308-384 per data point) and DiOHF (n=305-363) in NRVMs against pacing-induced CaMKII activity. **(G)** Cell viability after 48 hour compound incubation. All measurements are from biological triplicate studies. ns=p>0.05, ****p<0.0001; significance determined via one-way ANOVA and Dunnett’s multiple comparisons test (*C, E*).

We then evaluated whether ruxolitinib could replicate its CaMKII inhibition in vivo. 60-90 mg/kg dosing in mice is considered to be equivalent to the prescribed 20-25 mg doses in humans *(51, 52)*. 10 minute systemic pre-treatment with ruxolitinib at 41, 75, and 180 mg/kg suppressed isoproterenol-induced phosphorylation of phospholamban at threonine 17—a validated marker of CaMKII activity *(53–55)*—in a concentration-dependent manner (Fig. 4A-C). Altogether, our data demonstrate that ruxolitinib inhibits CaMKII in cardiomyocytes at clinically attainable concentrations, is well tolerated, and functions rapidly in vivo.

**Figure 4.**
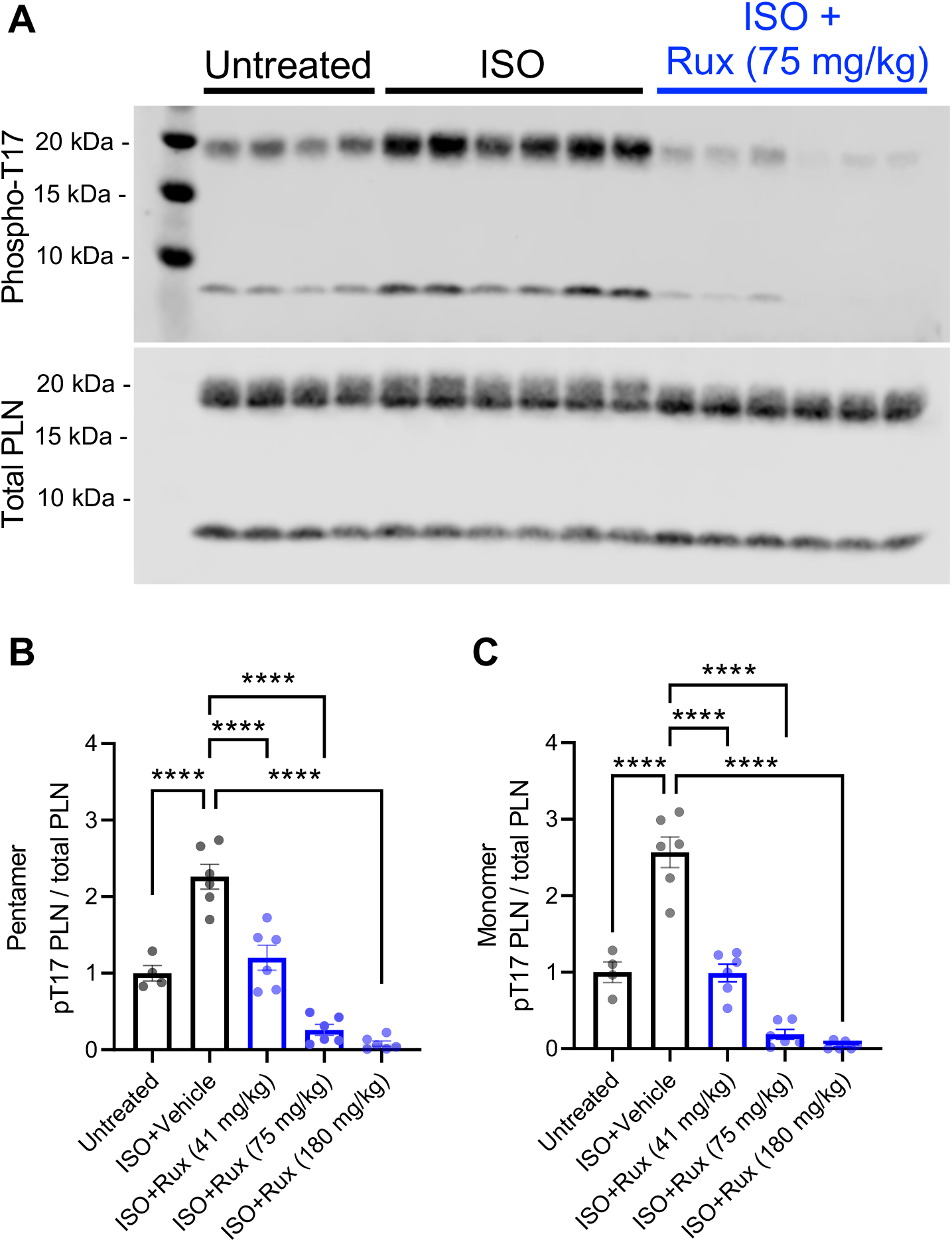
Ruxolitinib inhibits CaMKII in vivo. **(A)** Immunoblot and **(B to C)** quantitation of phospholamban (PLN) and Threonine-17 phosphorylated phospholamban in whole heart lysates from mice treated with intraperitoneal ruxolitinib for 10 minutes prior to isoproterenol stimulation. Data points represent individual mice. ****p<0.0001; significance determined via one-way ANOVA and Tukey’s multiple comparisons test.

### Ruxolitinib inhibits atrial fibrillation in hyperglycemic mice

Given its robust inhibition of CaMKII in cellulo and in vivo, we next tested whether ruxolitinib can ameliorate CaMKII-associated cardiac pathology. We chose atrial fibrillation as our focus since it is a major source of morbidity and mortality, affects nearly 50 million people worldwide globally, and is responsible for 20-30% of all ischemic stroke cases *(56, 57)*. CaMKII is a pivotal pro-arrhythmic signal in atrial fibrillation *(58–62)*. We previously validated a mouse model of atrial fibrillation enhanced by diabetes, a major risk factor for atrial fibrillation in patients and known upstream activator of CaMKII *(56, 62–65)*. In this model, genetic and chemical interventions that reduced CaMKII activity suppressed atrial fibrillation in diabetic mice *(62)*. 10-minute systemic pretreatment with 75 mg/kg ruxolitinib in hyperglycemic mice abolished pacing-induced atrial fibrillation (Fig. 5A and B). Examination of phosphorylated phospholamban in atria from these mice again confirmed suppression of CaMKII activity after ruxolitinib treatment (Fig. S7). Altogether, our data support that ruxolitinib can effectively inhibit CaMKII in both atrial and ventricular myocardium, and it can reverse or prevent arrhythmogenesis in experimental models acquired arrhythmia.

**Figure 5.**
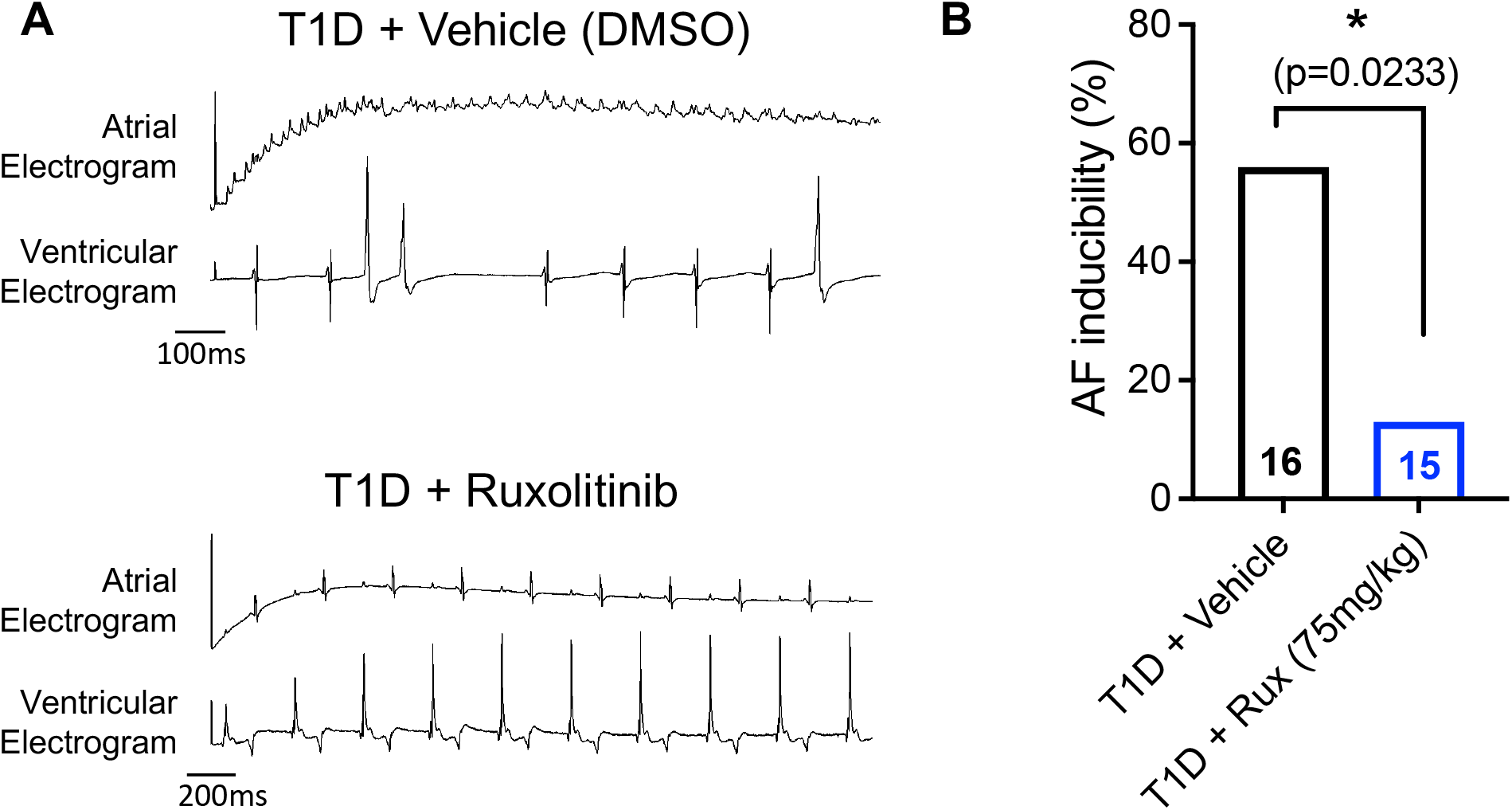
Ruxolitinib rescues atrial fibrillation in T1D mice. **(A)** Representative mouse atrial and ventricular tracings immediately after atrial burst pacing. DMSO treated mice demonstrate atrial fibrillation (upper panel), whereas 75 mg/kg ruxolitinib treated mice (single dose 10 minutes prior, intraperitoneal) retain sinus rhythm (lower panel). **(B)** Atrial fibrillation (AF) inducibility percentage from mice in A. Statistical comparison was performed using 2-tailed Fischer’s exact test.

### Ruxolitinib does not lead to short term or spatial memory deficits

CaMKII has long been known for its key role in learning and memory *(66)*. Thus, cognitive off-target effects have been a major concern and criticism toward therapeutic CaMKII inhibition *(29)*. Ruxolitinib is inefficient at crossing the blood-brain-barrier, with brain tissue reaching only 3.5% of plasma concentration in rats *(67)*. Furthermore, ruxolitinib has been prescribed to patients for over a decade and with no reported overt cognitive deficits *(68)*. This led us to hypothesize that there is a therapeutic window where efficient cardiac inhibition can be achieved without impairing learning and memory. To test if this was the case in our hands, we treated mice with ruxolitinib and subjected them to the novel object recognition test (NORT) and the Y maze spatial memory test (Fig. 6A)—both established behavioral paradigms that test short term and spatial memory. These tests were chosen because CaMKII inhibition is known to affect both short term and spatial memory *(69–71)*. Importantly, myriad studies observe robust deficits in novel object recognition after CaMKII inhibition *(69, 72–75)*. Neither single dose (1 hour prior) nor multiple doses (7-days, twice daily) of ruxolitinib (75 or 41 mg/kg doses) caused significant differences in novel object recognition, as measured by percent time spent with the novel object (Fig. 6B and C). Total distance traveled during testing was similar for both groups, confirming equivalent locomotor ability (Fig. 6D and E). In the Y maze test, single dosing using the lower dose showed decreased preference for the novel arm (Fig. 6F), but this effect was not seen in the higher dose or in either of the multiple dosing cohorts (Fig. 6F and G). Thus, we conclude that, at the tested doses, ruxolitinib does not lead to gross impairment of short term or spatial memory. These results demonstrate feasibility of cardiac CaMKII blockade without memory impairment in mice by using a potent inhibitor with inefficient uptake in the brain.

**Figure 6.**
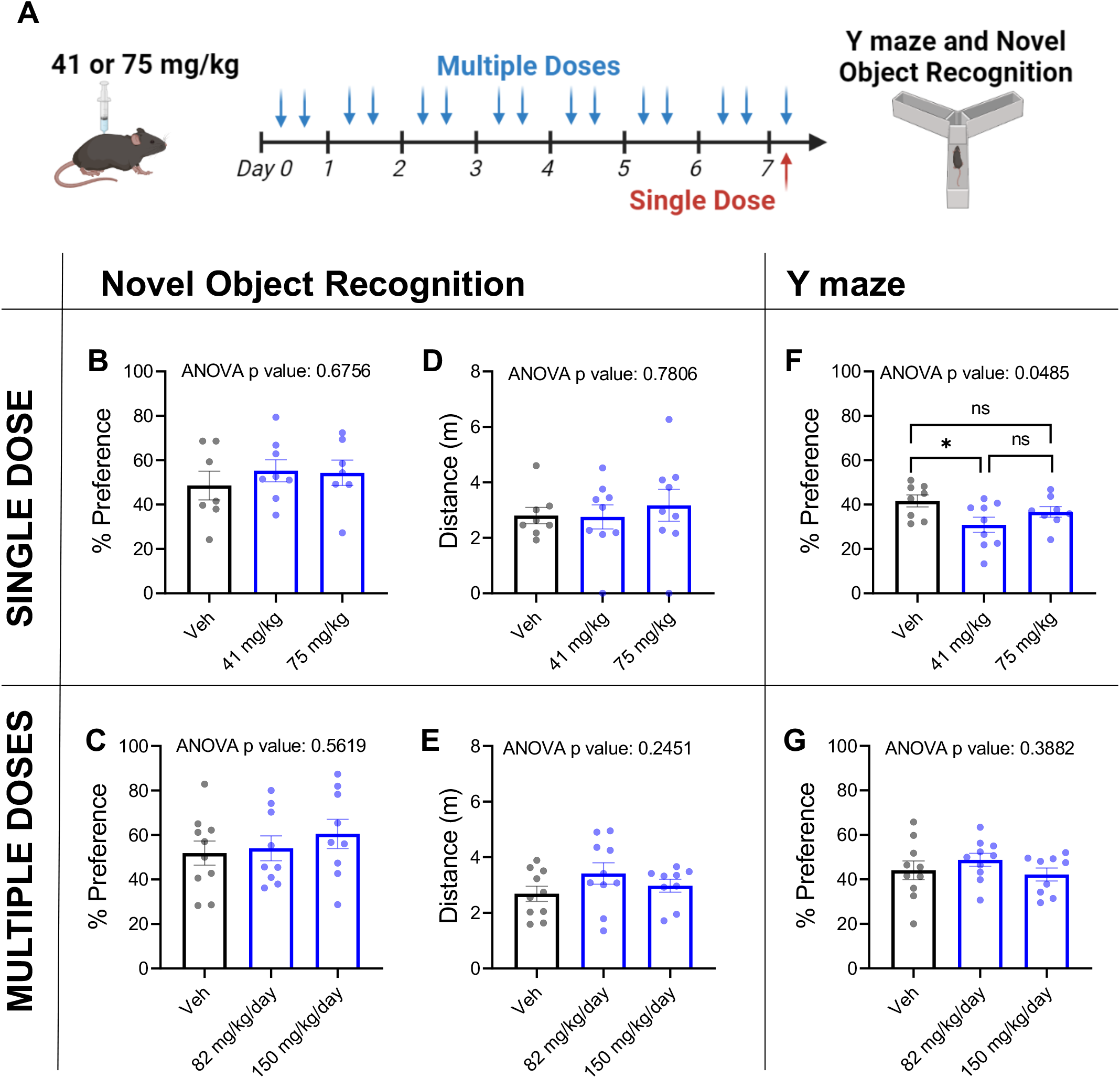
Ruxolitinib does not impair short term or spatial memory in mice. **(A)** Experimental paradigm: Wild type C57BL6J mice were treated with ruxolitinib by intraperitoneal injection acutely (1 hour) or chronically (7 days, twice daily injections) prior to cognitive testing via the novel object recognition test (NORT) and the Y maze spatial recognition memory test. **(B, C)** percent of time spent with novel object. **(D, E)** Total distance traveled while in testing chamber. **(F, G)** Percent of time spent in novel arm of Y maze. Each data point represents a mouse, n=9-10 mice for all conditions. ns=p>0.05, *p<0.05; significance determined by one-way ANOVA and Tukey’s multiple comparisons test.

## DISCUSSION

In this work, we developed and validated CaMKAR—a fluorescent CaMKII activity biosensor—and used it to identify ruxolitinib as a CaMKII inhibitor with substantial cardiotherapeutic potential. CaMKAR displays the highest sensitivity and dynamic range out of all CaMKII sensors to date; it enjoys a high degree of specificity and superior versatility afforded by ratiometric, intensiometric, and in vitro functionality. Thus, CaMKAR is uniquely suitable for high-throughput in cellulo screening, which simultaneously assesses cell permeability, toxicity, and potency toward CaMKII. We demonstrated these properties by screening a safe-in-human drug collection for CaMKII inhibitors. In our screen, the orally bioavailable, FDA-approved ruxolitinib emerged as a prime candidate for repurposing due to its potency and low cytotoxicity.

Ruxolitinib inhibits CaMKII in cardiac cells at clinically achievable concentrations both in cellulo and in vivo. While a 90 mg/kg dose in mice is equivalent to the human maximal prescribed dose (25 mg), 41 mg/kg was sufficient to ameliorate CaMKII activity and 75 mg/kg rescued cardiac arrhythmias. Thus, we are optimistic that CaMKII inhibition is feasible in humans using existing dosing. To demonstrate translational anti-arrhythmic potential we focused on atrial fibrillation, a common arrhythmia driven by CaMKII hyperactivity *(58–62)*. In a validated model of hyperglycemia-induced atrial fibrillation, ruxolitinib displayed near complete rescue of the arrhythmic phenotype. We are hopeful that therapeutic potential in this model can rapidly translate since the CaMKII-RyR2-dependent mechanism in atrial fibrillation is shared by numerous arrhythmias, both inherited—including catecholaminergic polymorphic ventricular tachycardia (CPVT) *(15, 76–79)*, Timothy and Barth Syndromes *(80, 81)*, Duchenne’s muscular dystrophy *(82)*, and Ankyrin B mutations *(83)*—and acquired—such as glycoside toxicity *(84)*, heart failure *(77)*, and alcoholic cardiomyopathy *(85)*.

Beyond therapeutic repurposing, our results add nuance to whether CaMKII inhibition is safe and feasible. While multiple studies have demonstrated that CaMKII blockade is cardioprotective, pharmaceutical companies have remained cautious about developing CaMKII inhibitors. Chief among their concerns is the pivotal role that CaMKII serves in memory and cognition *(66)*. Our results suggest that several drugs already in circulation, taken by thousands-to-millions of patients, inhibit CaMKII at clinically relevant concentrations. Cognitive testing in mice treated with ruxolitinib showed no overt memory deficit at doses that displayed robust cardiac CaMKII blockade. This is likely aided by the fact that ruxolitinib displays poor blood-brain-barrier penetrance with a ~29-fold higher concentration in plasma than brain in rats *(67)*, and is consistent with the lack of known cognitive side effects in humans. Thus, our results support a reinvigoration of CaMKII inhibitor development and show proof-of-concept that cardiac CaMKII blockade without impairing cognition is achievable with small molecules.

While our data suggest that ruxolitinib is a prime candidate for cardiac repurposing, excitement for such therapies must be tempered with a careful evaluation of ruxolitinib’s on-target effects. Systemic ruxolitinib is used to treat polycythemia vera, myelofibrosis, and splenomegaly. Due to on-target JAK1/2 inhibition, prolonged treatment can result in thrombocytopenia and anemia. Thus, we predict that repurposing will be most efficacious for indications that require rapid CaMKII inhibition, such as arrhythmias *(86, 87)*, ‘electrical storm’ *(88, 89)*, and acute ischemia/reperfusion injury *(6, 14, 19, 90–92)*. These indications are supported by ruxolitinib’s fast speed of action: treatment for just 10 minutes was sufficient to suppress catecholaminergic stress in vivo. These short treatment courses may offer an optimal tradeoff between CaMKII inhibition and on-target actions. Other experimental CaMKII-modifying modalities, such as anti-sense oligonucleotides and inhibitor-encoding gene therapy take effect in the order of days-to-weeks, which lessens their utility in these acute indications.

This study hints at exciting future directions. The unique collection of CaMKAR properties will enable addressing previously intractable questions. High sensitivity and fast kinetics will permit determining the spatiotemporal dynamics of cardiac CaMKII. This is an exciting area since recent work shows that CaMKII can be activated in a subcellular-dependent manner with differing effects and targets. Furthermore, CaMKAR can be extensively scaled in cell culture, which will permit more ambitious chemical screens to identify additional inhibitors. The molecules identified here can serve as scaffolds for medicinal chemistry in order to amplify their CaMKII potency while minimizing their originally intended effects. CaMKII is known to drive an extensive number of cardiac pathologies, and beyond its effects in myocardium, CaMKII has been shown to underpin other illnesses including asthma, metabolic disease, and cancer *(93– 96)*. Thus, a careful examination of the utility of these compounds in each of these disease-contexts is warranted.

## MATERIALS AND METHODS

### Study Design

The goal of this study was to engineer a sensitive and specific CaMKII biosensor suitable for high throughput screening. Once this was achieved, we sought to identify drugs that can be repurposed as CaMKII inhibitors. All animal studies were approved by the appropriate animal care and use committees. Mice of both sexes were used unless noted in the methods. Animal number was determined by prior experience and published reports. Mice were randomly assigned to treatment conditions, and importantly, whenever an experiment spanned across multiple days or housing cages, we ensured that all conditions were represented in these units to avoid batch effect confounders. Behavioral tests were performed by an independent experimenter blinded to the experimental conditions. Imaging data analysis was done in a computer-automated fashion to avoid human bias.

### Plasmids and Molecular Biology

To develop CaMKAR, we amplified cpGFP and FHA1 from pcDNA3.1(+)-ExRai-AKAR2 (gift from Jin Zhang, Addgene plasmid # 161753; http://n2t.net/addgene:161753; RRID:Addgene_161753) into pcDNA3.1 and pET-6xHis/TEV (VectorBuilder) plasmid backbones. CaMKII substrates were added to the 5’ end of cpGFP by site-directed mutagenesis (KDL Enzyme Mix, NEB). deadCaMKAR^T6A^ was generated by site-directed mutagenesis of pcDNA3.1:CMV-CaMKAR. For lentiviral delivery of CaMKAR and CaMKIIδ, CaMKAR was cloned into pLV plasmid backbone driven by hEF1a promoter to create pLV:hEF1a-CaMKAR-P2A-BlastR (VectorBuilder); CaMKIIδ was cloned into pLVX:TetONE-Puro-hAXL (gift from Kenneth Pienta, Addgene plasmid # 124797; http://n2t.net/addgene:124797; RRID:Addgene_124797). PKC sensor pcDNA3-ExRai-CKAR was a gift from Jin Zhang (Addgene plasmid # 118409 ; http://n2t.net/addgene:118409 ; RRID:Addgene_118409). Constitutively active CaMKI (pTwistCMV-CAMK1^T177D^) was synthesized by Twist Bioscience. Constitutively active CaMKIV-dCT was a gift from Douglas Black (Addgene plasmid #126422; http://n2t.net/addgene:126422; RRID:Addgene_126422). pCMV-Camui-NR3 was a gift from Dr. Michael Lin.

Purified CaMKAR was generated by transforming NEBExpress Competent E. coli (NEB) with pET-6Xhis/TEV-CaMKAR. After addition of Isopropyl Thiogalactoside (IPTG, 0.4 mM) and incubation at 15°C for 24 hours, cells were lysed and sonicated. CaMKAR was isolated from soluble lysate by nickel chromatography NEBExpress columns (NEB). Soluble protein was quantified by Pierce BCA assay (ThermoFisher) and stored in 50% glycerol at -80 °C.

### Cell Culture

#### Maintenance

HEK293T/17 cells (ATCC CRL-11268) were maintained in DMEM (L-glutamine, Sodium Pyruvate, Non-essential amino acids; Gibco) supplemented with 10% FBS (Gibco) and Pen/Strep (Gibco). Cells were maintained between 10%-95% confluence. K562 cells (ATCC CCL-243) were maintained in DMEM (Thermo Fisher) supplemented with 10% FBS and Penicillin/Streptomycin (Gibco). K562 cells were kept in suspension at a density between 100k-1M cells/mL. When confluent, cells were directly diluted into fresh media. NRVMs were isolated from Sprague-Dawley rats as previously described (XX). Isolated myocytes were cultured in DMEM (Gibco) supplemented with 10% FBS (Gibco) and Pen/Strep (Gibco). All cells were maintained in a humidified incubator at 37 ° with 5% CO_2_. To determine viability, cells were lysed in 1:1 PBS:CellTiter-Glo 2.0 (Promega) reagent. Lysate luminescence was assayed with a Synergy MX microplate reader.

#### Lentivirus production and infection

HEK293T/17 cells under passage #6 were seeded into 10cm dishes at 400k cells/mL. These cells were transfected using TransIT-Lenti (Mirus Bio) using a ratio of 5:3.75:1.25 of packaging plasmid:psPAX2:pMD2.G for a total of 10 µg per dish. After 48 hours, supernatant was collected, clarified, and concentrated 10-fold using Lenti-X concentrator (Takara). Lentivirus aliquots were stored at -80 °C until functional titration and use. Infection occurred at indicated multiplicity of infection in the presence of 10 μg/mL polybrene reagent (Sigma).

#### Plasmid Transfection

HEK293T/17 cells were plated into well plates at 400k cells/mL unto PDL coated 24 well plates. Each well was transfected with 500 ng of DNA complexed with 1 µL of JetPrime reagent. Cells were examined 24-48 hrs post transfection.

#### Adenovirus infection

CMV-CaMKAR-encoding Ad5 adenovirus was synthesized directly by Vector Biolabs. NRVMs were infected 24 hours post isolation at multiplicity of infection 100. Cells were analyzed 48 hours post infection.

### Microscopy

Timelapse microscopy was performed using an Olympus IX-83 inverted widefield microscope equipped with an ORCA Flash 4.0 and Lumencor SOLA light source. CaMKAR signal was captured at 200 ms exposure using the following channels: excitation filters ET402/15x and ET490/20x and emission filter ET525/35m (Chroma Technology). CaMKAR Signal (R) is defined as the ~488-nm-excited intensity divided by the ~400-nm-excited intensity. Calcium imaging was collected in the TRITC channel ex 555/em. 590 nm. Confocal imaging was performed using a Zeiss LSM880 Airyscan FAST. Using excitation lasers 405 nm and 488 nm and collecting emission window at 520 ± 10 nm.

#### Image analysis

Otsu segmentation was used track individual cells and their intensity in 488 nm and 400 nm channels at every timepoint in CellProfiler. Individual cell and well values were imported to R Studio for tabulation and summary statistics including mean, standard deviation, and n calculation.

### CaMKAR-based high throughput screening

We created a polyclonal K562^CaMKAR-CaMKII^ line by infecting 3 million cells with lentiviruses containing pLVX:EF1a-CaMKAR-BlastR (multiplicity of infection=5) and pLVX:TetONE-CaMKIIT287D-P2A-mCherry (multiplicity of infection=1). These cells were expanded to 600M and incubated with 1 µg/mL Doxycycline 24 hours prior to screening. Cells were monitored for mCherry expression to confirm CaMKII expression. All cells were pelleted and resuspended in Live Imaging Cell Solution (Gibco) supplemented with 1 µg/mL Doxycycline and 4.5 g/L glucose. These cells were then distributed across 15 384 clear-bottom well plates at 50k cells per well. 5 µL of 10x compound from the Johns Hopkins Drug Library v3.0 was added at a final concentration of 5 µM per compound. The Johns Hopkins Drug Library v3 was assembled by combining the Selleckchem FDA-Approved Drug Library (3,174 compounds, Catalog No: L1300) with non-duplicate compounds found in APExBIO DiscoveryProbe FDA-approved Drug Library (483 compounds, Catalog No. L1021) and the MicroSource US Drug Collection (817 compounds). The library was formatted for use in 384-well plates and stored in DMSO at -80 °C. Each plate contained 2 columns of untreated cells, and 1 column of cells treated with AS100397 10 μm. Cells were incubated for 12 hours before reading by high content imager (MolDev IXM High Content Imager). Images were automatically segmented and quantified using CellProfiler as above. R studio was used to quantified the ratio between 488 nm and 405 nm excitation channels as above. CaMKII inhibitory % was calculated by min-max normalizing the data between untreated and AS100397-treated samples. 118 candidates were selected as those with 60% inhibition or better, which represents a false discovery rate adjusted p-value (Benjamini-Hochberg) cutoff of <0.003.

#### In vitro subscreen

118 candidates from in cellulo screen were then subjected in vitro CaMKAR screen. In vitro CaMKAR reaction was carried in 384-well plates with 25 μL total volume, 12.5 μL 2X Kinase Buffer (), 1.25 μL 10mM CaCl, 0.5uL 50 μM CaM, 0.11 μL CaMKAR (25 pmol), 0.0625 μL CaMKII (125 fmol), completed with nuclease-free water. Plate was read at baseline via Tecan Safire microplate reader, then 5 μL of 5x drug was added (5 μM final), plate read again, then 5 μL ATP was added (100 μM final) and kinetic assay was read for 60 mins. Slope in the change of CaMKAR ratio was used as metric for CaMKII activity. Data was min-max normalized between untreated and AS100397-treated wells. Hits were defined by significant deviation from untreated distribution by the z-Statistic p-Value. This statistical approach was reviewed by the Johns Hopkins Biostatistics, Epidemiology, and Data Management Core.

### Mouse models

All mouse studies were carried out in accordance with guidelines and approval of the Johns Hopkins University Animal Care and Use Committee (Protocol #MO20M274). Male C57BL/6J mice (The Jackson Laboratory, ME, USA) were housed in a facility with 12 hr light/12 hr dark cycle at 22 ± 1 °C and 40 ± 10% humidity. Teklad global 18% protein rodent diet and tap water were provided ad libitum.

#### Isoproterenol challenge

ruxolitinib phosphate (MedChemExpress) was injected via intraperitoneal injection. 10 minutes later, 5 mg/kg isoproterenol (Sigma-Aldrich) was injected via intraperitoneal injection. Mice were euthanized 10 minutes after the isoproterenol challenge.

#### T1D Mouse Model

Type-1 diabetes (T1D) was induced as previously described *(62)*, in adult (8 – 12-week-old) male mice C57BL/6J mice (The Jackson Laboratory, ME, USA) by a single intraperitoneal (i.p.) injection of streptozotocin (STZ) (185mg/kg, Sigma-Aldrich) dissolved in a citrate buffer (citric acid and sodium citrate, pH 4.0) after a six hour fast. Mice were maintained on normal chow diet (NCD) (7913 irradiated NIH-31 modified 6% mouse/rat diet – 15 kg, Envigo, Indianapolis, IN). Mice were considered diabetic if blood glucose level was ≥ 300mg/dl via tail vein blood checked with a glucometer -OneTouch Ultra 2 (LifeScan, Zug, Switzerland). Invasive electrophysiology study was done 2 weeks after STZ injection.

#### AF induction

Anesthesized mice were injected via i.p. route with either Ruxolitinib phosphate (MedChemExpress) – 75mg/kg dissolved in 10% Dimethylsulfoxide (DMSO) or vehicle (10% DMSO) 10 minutes prior to rapid atrial burst pacing to assess atrial fibrillation (AF) inducibility. In vivo electrophysiology (EP) studies were performed as previously reported *(62)* in mice anesthetized with isoflurane (2% for induction and 2 % for maintenance of anesthesia; Isotec 100 Series Isoflurane Vaporizer; Harvard Apparatus). AF was defined as the occurrence of rapid and fragmented atrial electrograms with irregular AV-nodal conduction and ventricular rhythm for at least 1 second. If 1 or more atrial bursts (out of 5) were an AF episode, the mouse was considered to have inducible AF. Mice were euthanized immediately following the procedure and atrial tissue from both right and left atria were obtained and flash frozen in liquid nitrogen and then stored at -80°F.

### Western Blotting

Mouse hearts were disrupted using a tissue blender in 1% Triton X-100 containing protease and phosphatase inhibitors. To detect phospho- and total phospholamban, samples were left unboiled to preserve pentameric form. Samples were run on 4-12% bis-tris acrylamide gels and transferred unto nitrocellulose membranes. Total and phosphorylated-T17 phospholamban were assayed by incubation with respective antibodies (Phospholamban pT17 pAb (Badrilla A010-13), 1:5000; Phospholamban (Thermofisher/Pierce MA3-922), 1:2000) followed by secondary labeling (Goat Anti-Rabbit IgG Antibody 680LT (Licor 926-68021) 1:10,000; Goat Anti Mouse IgG Antibody 800CW (Licor 926-32210) 1:10,000) and quantified using a LICOR Odyssey imager and quantified with ImageStudio.

### Behavioral paradigms

Memory tests were performed at the Johns Hopkins Behavioral Core by a blinded experimentalist. Short term memory was assessed using the novel object recognition test. The test consisted of three phases over two days. On day 1 mice were habituated to the arena for 10 minutes. On day 2, mice were allowed to explore the arena consisting of two identical objects, referred to as the familiar objects, for 10 minutes and were then placed back in the home cage for 30 minutes. Following the delay, the mice were placed back into the arena with one “familiar” object and one “novel” object and allowed to explore the objects for 5 minutes. Distance travelled and time spent investigating each object was automatically recorded using Anymaze tracking software (Stoelting Co., Wood Dale, IL, USA).

Spatial memory was tested via the Y-maze spatial recognition test. The Y-maze consists of three 38 cm-long arms (San Diego Instruments). During the training phase, one arm of the Y-maze was blocked. The mouse was placed at the end of one of the two open arms and allowed to explore for 5 min. After a 30-min inter-trial interval, the test phase began: the blockade was removed, and the mouse was allowed to explore all three arms of the maze for 5 min. Distance traveled and time spent in each arm was automatically recorded using Anymaze tracking software. Data from the first 2 min of the test phase were used to evaluate percent time spent in the novel arm.

### Statistics and schematics

Imaging summary data condensed and organized with R Studio. Statistical testing done with GraphPad Prism v8.2.0 as described in each figure legend. Schematics and drawings made with Biorender.com. CaMKAR Ratio pseudocoloring for fluorescence microscopy performed with ImageJ using the Ratio Plus plugin. Kinase dendrogram illustration made with KinMap *(97)* and reproduced courtesy of Cell Signaling Technology, Inc. (www.cellsignal.com). Signal-to-noise ratio was calculated as previously described *(41)*: single-cell maximal ratio change after stimulation divided by the standard deviation across 4 baseline timepoints.

## Supporting information

Supplemental datafile 1

Supplemental datafile 2

## List of Supplementary Materials

Supplementary Materials and Figures S1-S8 Data files S1-S2

## Acknowledgments

We are thankful to the Johns Hopkins Biostatistics, Epidemiology, and Data Management Core for statistical analysis stewardship, to Dr. David A. Kass for providing neonatal rat ventricular myocytes and pacing system and to Fondation Leducq for providing AS100397 compound.

## Funding

American Heart Association Predoctoral Fellowship 905878 (OERG)

National Institutes of Health R35 HL140034 to (MEA)

Sarnoff Fellowship to (LJN)

## Author contributions

Conceptualization: OERG, MEA

Methodology: OERG, AL, BLL, JOL

Investigation: OERG, JMG, LJN, CET, OOM, KLS, EDL

Visualization: OERG

Funding acquisition: OERG, LJN, MEA

Project administration: OERG, MEA

Supervision: OERG, MEA

Writing – original draft: OERG

Writing – review & editing: OERG, MEA

## Competing interests

The authors declare that they have no competing interests.

## Data and materials availability

All data associated with this study are present in the paper or the Supplementary Materials. Raw image files and AS100397 compound are available upon request.

## Notes

### Competing Interest Statement

The authors have declared no competing interest.

