## Supplemental Figures for "Novel Biosensor Identifies Ruxolitinib as a Potent and Cardioprotective CaMKII Inhibitor"

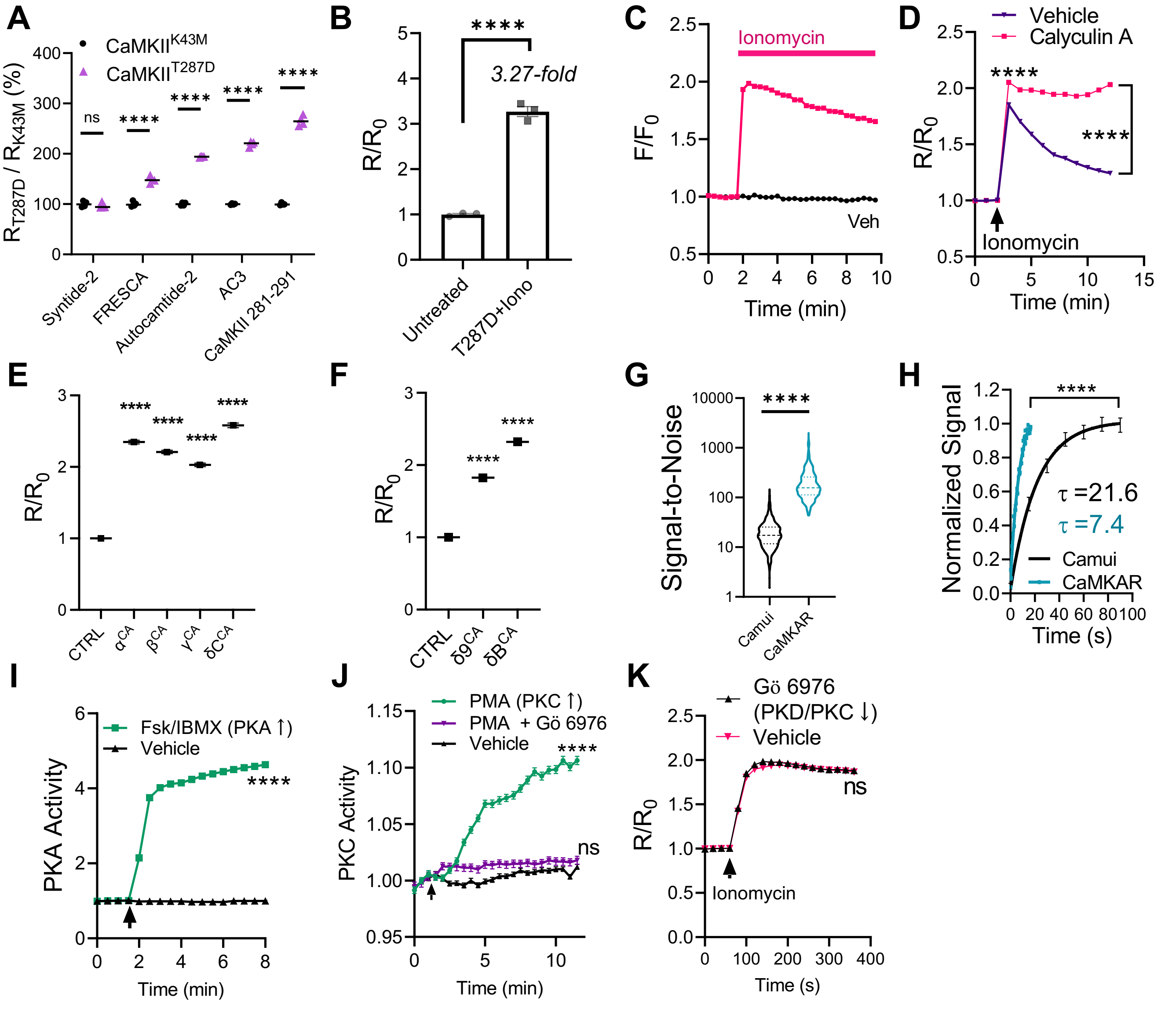


**Figure S1. CaMKAR development and validation. (A)** Screening to identify peptide that renders sensor sensitive to CaMKIIδ: plate reader fluorometry of 293T cells transfected with CaMKIIδ and prototype sensors. N=3 wells per observation. **(B)** Maximal stimulation of CaMKAR in 293T cells by expression of CaMKII$^{T287D}$ and ionomycin treatment (5 μM; n= mean of 3 wells) or untreated with either (n=mean of 3 wells). **(C)** Intensiometric CaMKAR signal (Ex. 488 nm, Em. 520 nm) after vehicle (Veh, n=2,488-2,615 cells) or ionomycin stimulation (n=1,733-2,220 cells), from same dataset as Fig. 1C. **(D)** CaMKAR signal in 293T cells pre-treated with vehicle (n=1,746-1,930 cells) or Calyculin A (10 nM; n=1,051-1,595), then stimulated with ionomycin. **(E)** CaMKAR response in control (CTRL; n=151) cells or after transfection of constitutively active CaMKII α (n=194 cells), β (n=229), δ (n=201), or γ (n=194) and **(F)** control (CTRL; n=772) vs constitutively active CaMKIIδ splice variants 9 (n=1,136) and B (n=1,140). **(G)** CaMKAR and Camui-NR3 signal response to ionomycin in 293T cells: assayed for signal-to-noise ratio (CaMKAR n=447 cells; Camui n=469 cells), and **(H)** activation kinetics (CaMKAR n=73 cells; Camui n=3,576-4,147 cells). **(I)** 293T cells expressing PKA sensor ExRai-AKAR2 treated with vehicle (n=1,066-1,092 cells) or Fsk (50 μM)/IBMX (100 μM)(n=908-1,135 cells). **(J)** 293T cells expressing PKC sensor ExRai-CKAR treated with vehicle (n=1,490-1,591 cells), PMA (100 ng/mL; n=932-970), or PMA + Gö 6976 (500 nM; n=770-823). **(K)** Effect of vehicle (n=722-792) or pre-treatment with Gö 6976 (500 nM; n=509-587) on ionomycin CaMKAR response. Arrows denote addition of compound. Data shown as mean ± SEM (unless too small to display). All observations done in biological triplicate. ns=p>0.05, ****p<0.0001; significance was determined via one-way ANOVA and Sidak's multiple comparisons test (*A*) or Dunnett's multiple comparisons test (*E and F*); two-way ANOVA and Sidak's multiple comparison test (*D, I, and K*) or Tukey's multiple comparisons test (*J*); unpaired two-tailed T-test (*B and G*); or nonlinear regression (*H*).

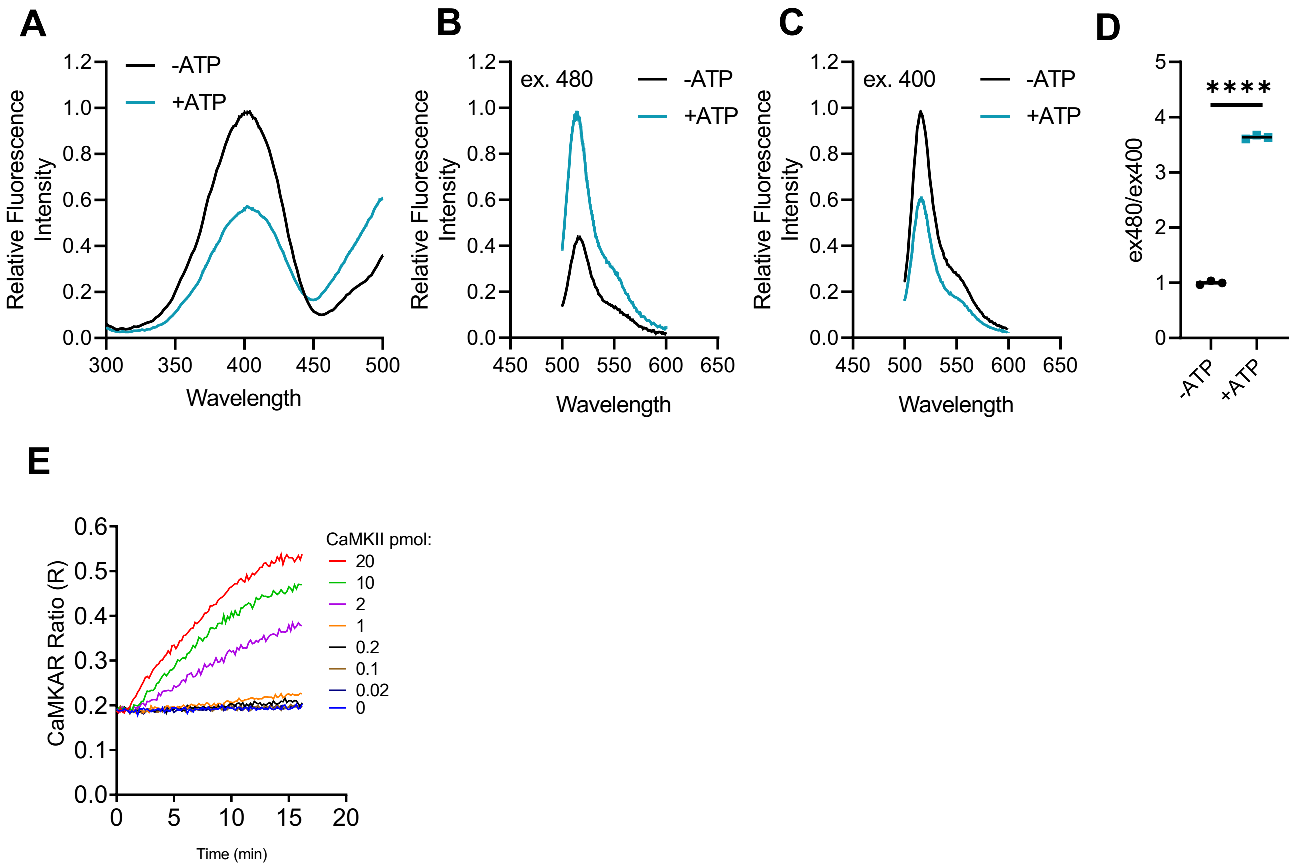


**Figure S2. CaMKAR is functional in vitro.** Purified recombinant CaMKAR was co-incubated with CaMKIIδ$_C$, $Ca^{2+}$/CaM ± ATP and assayed via fluorescence plate reader for **(A)** excitation spectrum at 520 nm emission, **(B)** emission spectrum at 400 nm excitation, **(C)** emission spectrum at 488 nm excitation, and **(D)** maximal CaMKAR ratio (see methods). **(E)** Purified CaMKAR co-incubated with increasing concentrations of purified CaMKIIδ$_C$ (see methods). Data shown as mean ± SEM (unless error is too small to graph). Observations done in technical triplicate (*A-D*) or single replicate (*E*).****=p<0.0001; significance determined via unpaired two-tailed T-test.

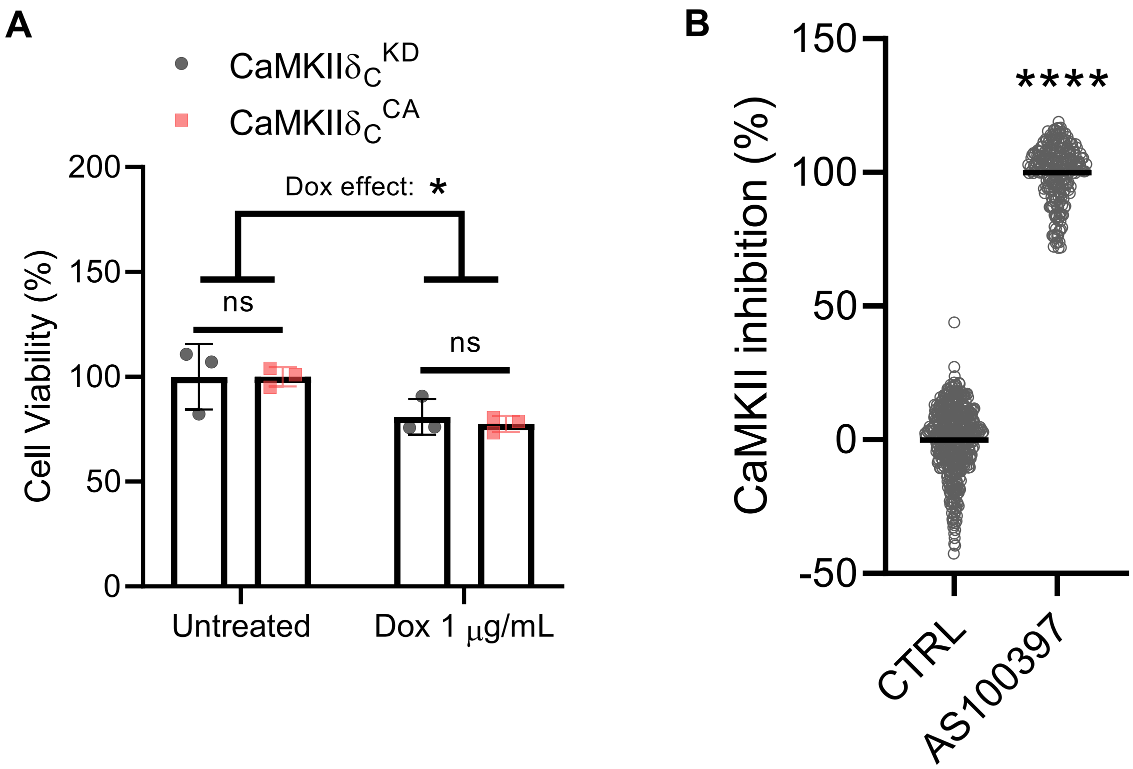


**Figure S3. K562 cells tolerate CaMKIIδ$^{T287D}$ overexpression and are suitable for screening.** K562 cells infected with lentiviruses encoding CMV-driven CaMKAR and TetON-driven kinase-dead or constitutively active CaMKIIδ$_C$ were treated with doxycycline for 24 hours and assayed for viability via CellTiterGlo 2.0 luminescence. Cells displayed doxycycline-associated toxicity but tolerated hyperactive CaMKII. Observations done in biological triplicate. Data points represent individual wells normalized to untreated condition. ns=p<0.05, *p<0.05; significance determined via two-way ANOVA and Tukey's multiple comparisons test. **(B)** CaMKAR signal in K562$^{CaMKAR/CaMKII}$ cells co-incubated with AS100397 (10 μM, n=240 wells) or vehicle (n=504 wells) in 384 well plate format. Data min-max normalized to 0-100% based on means of the two groups. ****p<0.0001; significance determined via unpaired Student's t -test.

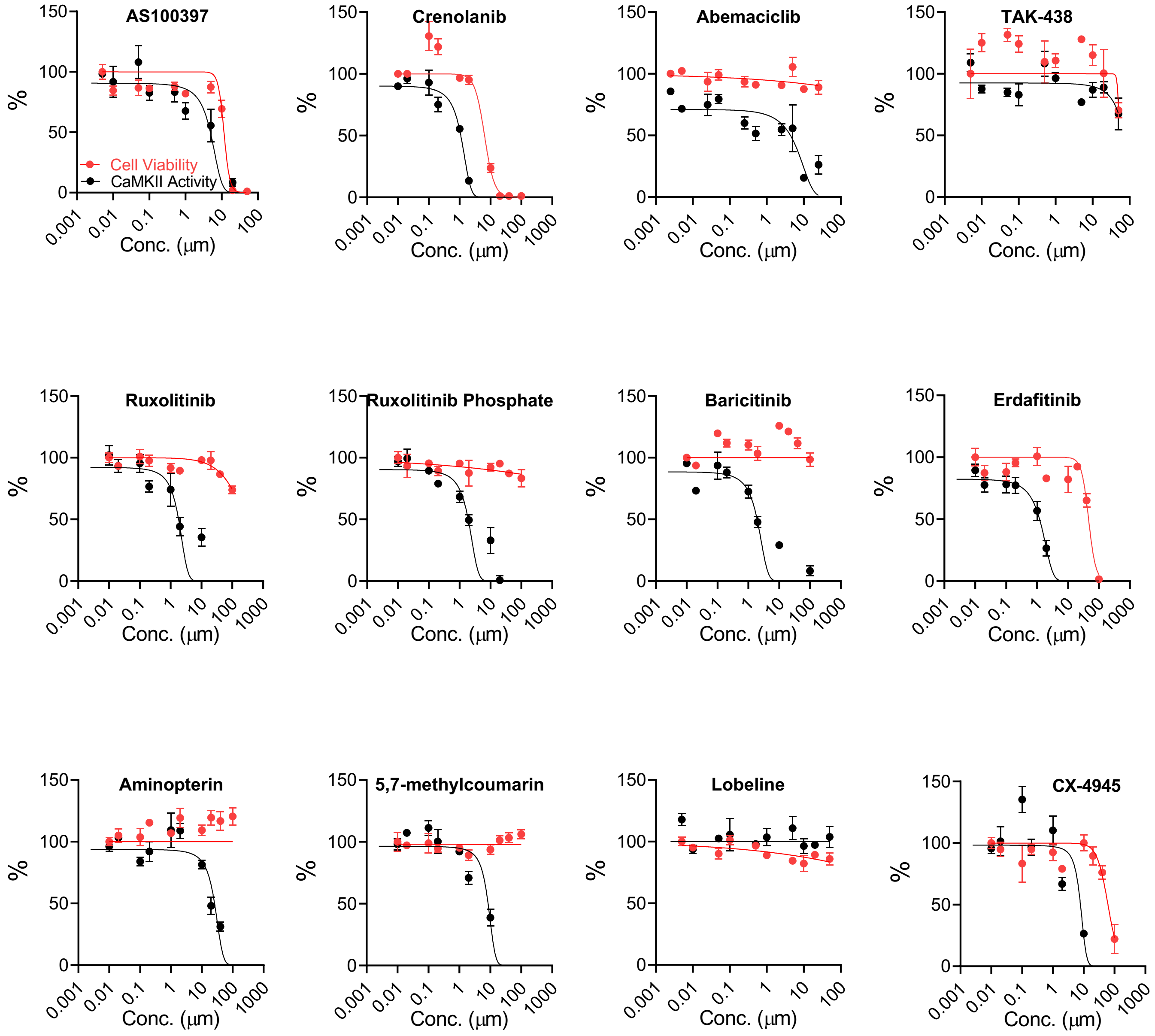


**Figure S4. CaMKII inhibition and cell viability in 293T cells**. CaMKAR- and CaMKII$^{T287D}$-expressing 293T cells were treated with hits identified in fig. 3C, AS100397 as positive control. Compounds were incubated for 12 hours and assayed via high content imaging for CaMKII activity and via CellTiter-Glo 2.0 luminescence for cell viability. Data points represent mean ± SEM from 3 independent wells each.

Ruxolitinib

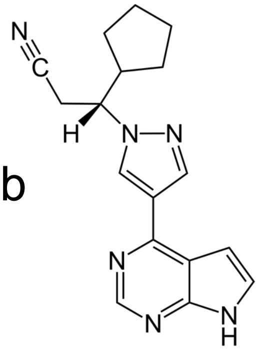

Crenolanib

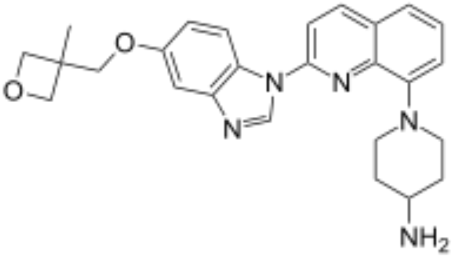

Baricitinib

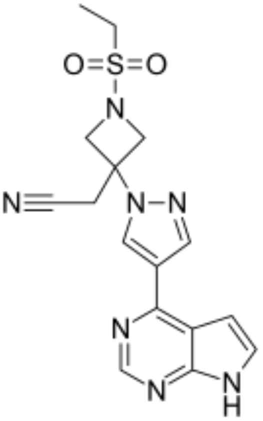

Abemaciclib

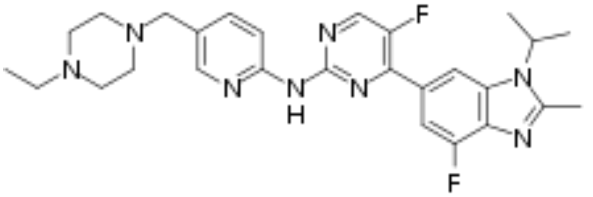

Silmitasertib

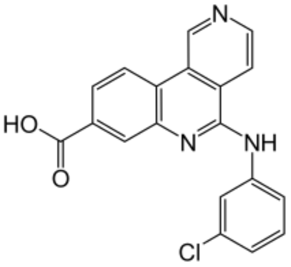

**Figure S5.** Chemical structures of identified CaMKII inhibitors.

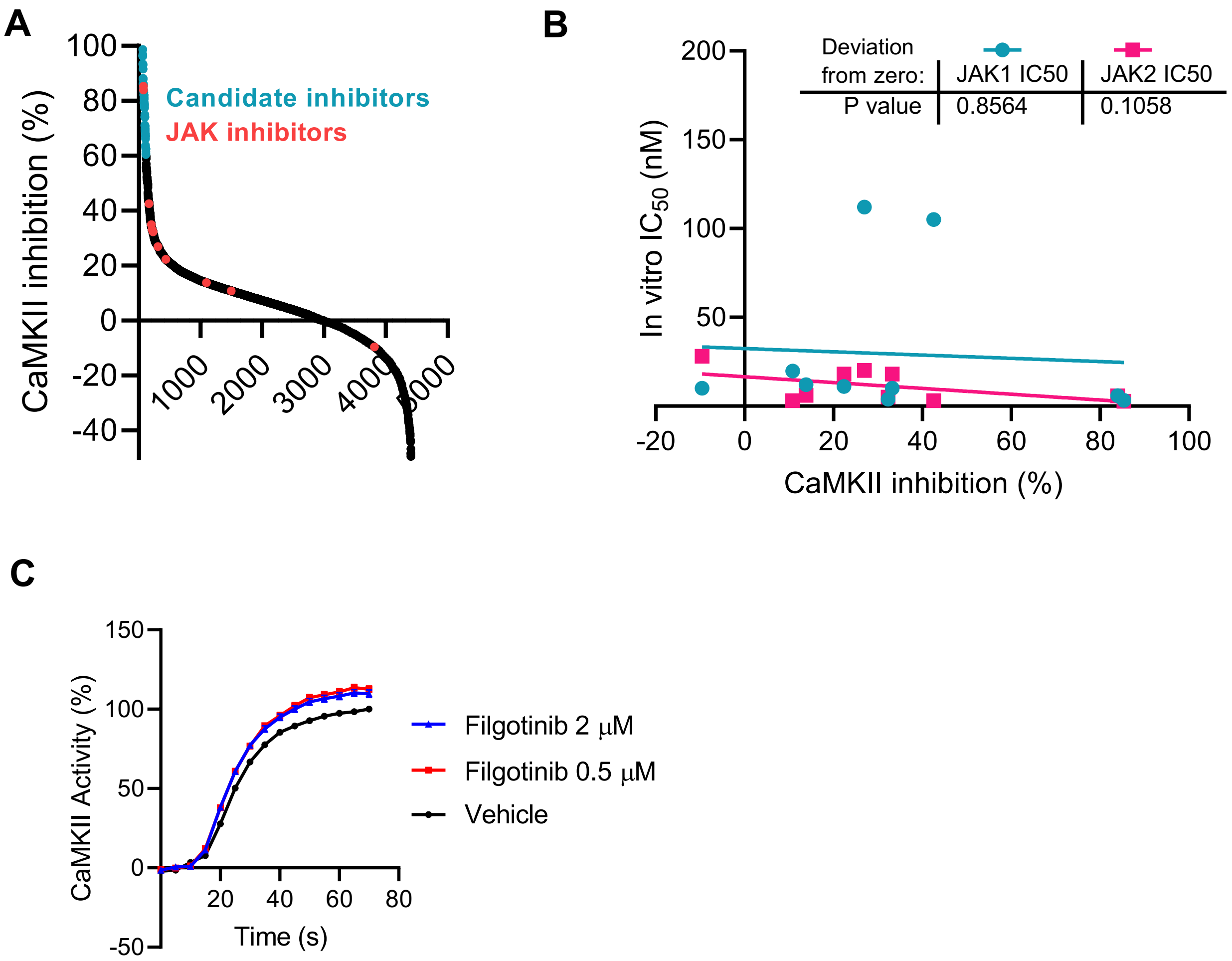


**Figure S6. CaMKII inhibition is independent of JAK1/2 inhibition. (A)** Known JAK1/2 inhibitors included in the primary screen (from Fig. 2b) are displayed in red. **(B)** JAK1/2 inhibitors from *A* plotted by their known in vitro $IC_{50}$ against JAK1 and JAK2 versus their CaMKII inhibition score. *Insert:* linear regression analysis to determine significance of correlation. **(C)** Pacing-induced CaMKII activity in CaMKAR-expressing NRVMs treated with vehicle (n=284-354) JAK inhibitor filgotinib (0.5 μM; n=291-335; 2 μm n=290-330 cells).

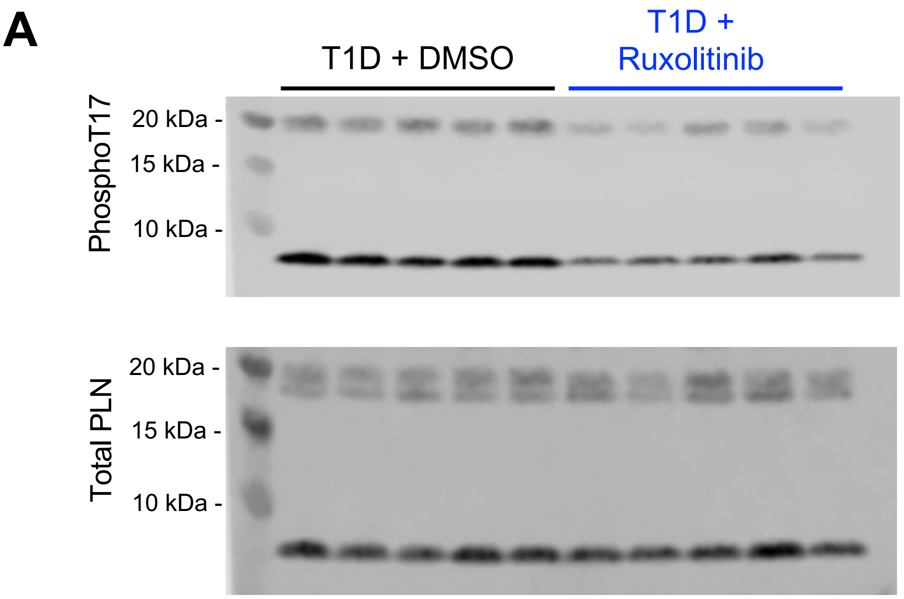


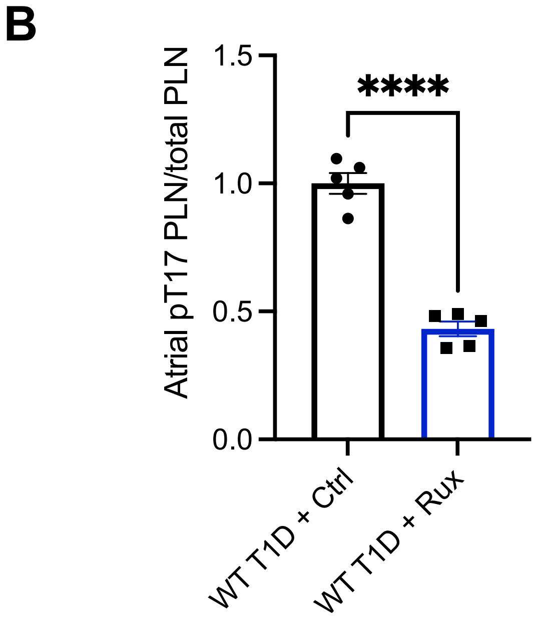


**Figure S7. Ruxolitinib inhibits CaMKII in atria. (A)** Atrial heart lysates prepared from T1D animals treated with ruxolitinib were separated by SDS-page electrophoresis and probed with specific antibodies to phospholamban (PLN), phosphorylated phospholamban (pPLN). **(B)** Quantification of immunoblot from A. Data points represent individual mice. Significance was determined via unpaired T-test; ****$p<0.0001$.
